# Convergence of two global regulators to coordinate expression of essential virulence determinants of *Mycobacterium tuberculosis*

**DOI:** 10.1101/2022.06.30.498252

**Authors:** Hina Khan, Partha Paul, Ritesh Rajesh Sevalkar, Sangita Kachhap, Balvinder Singh, Dibyendu Sarkar

**Author notes:** Address correspondence to: Dibyendu Sarkar, CSIR-Institute of Microbial Technology.

## Abstract

Cyclic AMP (cAMP) is known to function as a global regulator of *M. tuberculosis* gene expression. Sequence-based transcriptomic profiling identified the mycobacterial regulon controlled by the cAMP receptor protein, CRP. In this study, we identified a new subset of CRP-associated genes including virulence determinants which are also under the control of a major regulator, PhoP. Our results suggest that PhoP as a DNA binding transcription factor, impacts expression of these genes, and phosphorylated PhoP promotes CRP recruitment at the target promoters. Further, we uncover a distinct regulatory mechanism involving CRP and PhoP on transcriptional control of these genes. While both these regulators influence gene expression, we show that activation of these genes requires direct recruitment of both PhoP and CRP at their target promoters. The most fundamental biological insight is derived from the inhibition of CRP binding at the regulatory regions in a PhoP-deleted strain owing to CRP-PhoP protein-protein interactions. Based on these results, a model is proposed suggesting how CRP and PhoP function as co-activators and contribute to activation of the essential pathogenic determinants. Together, these results uncover a novel mechanism where two interacting virulence factors impact expression of virulence determinants, have significant implications on TB pathogenesis.

## Introduction

*Mycobacterium tuberculosis*, as one of the most successful human pathogens, retains the ability to adapt to diverse intracellular and extracellular environment it encounters during infection, persistence, and transmission [1]. Thus, *M. tuberculosis* is able to survive within interior of macrophages, and droplet nuclei in the atmosphere that are generated by individuals infected with the bacilli. While inhalation of droplets causes the disease spread, following infection *M. tuberculosis* can persist in a non-replicating (dormant) state for a long period of time. A characteristic feature of tuberculosis is that the disease actually occurs through reactivation of dormant infection under favourable conditions (for example, under compromised immunity of infected individuals). In keeping with this, about a third of the world’s population is believed to have latent *M. tuberculosis* infection [2] with a lifetime risk of reactivation of TB as high as 5-10% [3]. Therefore, appropriate regulation of gene expression in mycobacteria is considered critical for establishing and emerging from the dormant state.

More than 100 transcription regulators, 11 two-component systems, 6 serine-threonine protein kinases and 13 alternative sigma factors are present in *M. tuberculosis*, suggesting that the complex transcriptional regulation is important for mycobacterial pathogenesis. Thus, to understand *in vivo* physiology of *M. tuberculosis* it is becoming increasingly important to investigate the role of individual regulators and their participation in integrated networks. In the recent past, cAMP has been shown to function as a global regulator of mycobacterial gene expression with a critical involvement in host-pathogen interactions and virulence [4–6]. Importantly, *M. tuberculosis* genome comprises of 17 genes encoding adenyl cyclases [7] and it has been suggested that dynamic balance of cAMP synthesis and secretion plays a key role in infection [8]. Also, Rv0386 is one of these adenyl cyclases which is required for virulence [9]. Little is yet known about the regulation of cAMP-associated mycobacterial genes.

The classical model of bacterial cAMP regulation is best studied in *E. coli* [10], where cAMP signal is transduced by CRP. Upon cAMP binding, CRP undergoes a conformational change, shows an enhanced recognition to specific DNA sequence and plays a central role to coordinate global transcriptional regulation for optimal bacterial utilization of different carbon sources. High throughput ChIP sequencing data suggests that *E. coli* CRP shows extensive low-affinity binding, suggesting a possible role of the regulator as a chromosome organizer [11]. The corresponding *M. tuberculosis* CRP (encoded by Rv3676) [5, 12, 13], which shares a sequence identity of 32% and similarity of 53% over 189 residues of *E. coli* CRP [14], recognizes a similar DNA binding motif and is shown to regulate transcription of a large number of mycobacterial genes [14]. Although cAMP binds to *M. tuberculosis* CRP with weaker affinity and lesser impact on DNA-binding[15], polymorphisms in CRP which result in enhanced DNA binding have been reported to influence transcription of a number of genes for *M. bovis* BCG [16, 17]. Consistent with its global role in transcription regulation, a CRP-deleted mutant is implicated in virulence because of its significant growth attenuation in mice and macrophages [14]. Further, CRP shows the highest activation of *rpfA* and *whiBI* genes [14], encoding proteins that are involved in reviving dormant bacteria [18] and whiBI encodes wbI family proteins [19, 20] that function to control developmental processes. Thus, CRP is implicated in controlling mycobacterial developmental processes that regulates persistence and/or emergence from the dormant state. However, mechanisms of transcription activation of CRP-regulated mycobacterial genes in response to metabolic signals remains poorly understood.

While probing genome-wide transcriptomic profile coupled with ChIP-seq/SELEX data of major regulators, we observed that a subset of genes is under the positive regulation of two major regulators, CRP [21] and PhoP [22–25]. Thus, we sought to define scope of individual regulators and their participation in integrated regulatory networks. Here, we uncover a novel role of PhoP in the mechanism of CRP-dependent mycobacterial gene expression. While these results account for the failure to activate a few CRP-regulated genes in *ΔphoP*-H37Rv, we provide evidence showing that an activation of a subset of genes requires simultaneous presence of both CRP and PhoP at the target promoters. Insightfully, results reported in this work account for an explanation of how CRP-dependent activation of a subset of mycobacterial genes is also linked to positive regulation by PhoP [23, 26], underscoring a critical role of PhoP in virulence gene expression via a complex mechanism of transcriptional control involving functional cooperation of the two virulence regulators.

## RESULTS

### *phoP* directly regulates a subset of genes of CRP regulon

DNA microarray and sequence-based transcriptomic profiling identified PhoP regulon [23, 26]. The role of the virulence regulator PhoP in cAMP-responsive mycobacterial gene expression became apparent when we noted an overlap between the regulons under the control of cAMP receptor protein (CRP) and PhoP. Thus, we compared relative expression of representative CRP regulon genes in WT and *ΔphoP*-H37Rv under normal and indicated stress conditions (Fig. 1A). Our results demonstrate that a subset of CRP-regulated genes including *icl1*, *umaA* and *whiB1* are positively regulated by PhoP, suggesting that PhoP also functions as an activator of these genes. Notably, *icl1* has been implicated in persistence and virulence of *M. tuberculosis* in macrophages and mice [27–32], whereas *umaA* encodes for a mycolic acid synthase [33]. On the other hand, *whiB1* as an essential gene has been shown to function in NO sensing of mycobacteria [34]. Since we noted elevated expression of these CRP-regulated genes in the complemented mutant, we compared *phoP* expression in WT-H37Rv and complemented *ΔphoP*-H37Rv (Fig. S1). Our results demonstrate that *phoP* expression level is reproducibly higher in the complemented mutant (relative to the WT bacilli) both under normal conditions (Fig. S1A) as well as during growth under low pH conditions (Fig. S1B). These results possibly account for an elevated mRNA levels of a few representative genes in the complemented mutant relative to WT-H37Rv (Fig. 1A). However, relatively poor restoration of gene expression in the complemented mutant under acid stress is possibly related to inadequate activation of PhoPR under specific stress conditions.

**Fig. 1:**
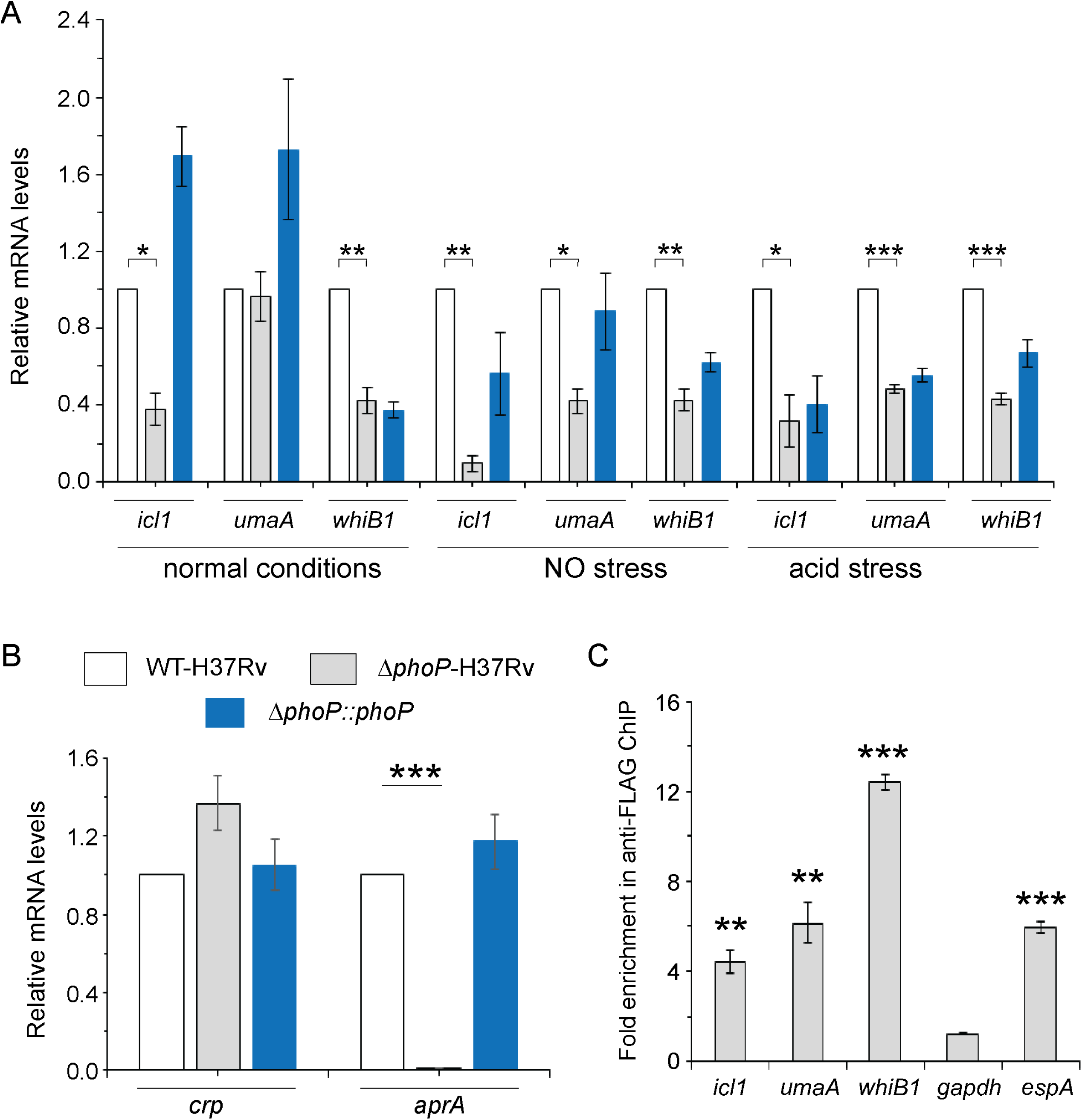
PhoP-depletion impacts expression of a subset of CRP-regulon genes. (A) Quantitative RT-PCR was carried out to compare expression level of *icl1*, *umaA* and *whiB1* in indicated mycobacterial strains, grown under normal and specific stress conditions. The results show average values from biological triplicates, each with two technical repeats (*P≤0.05; **P≤0.01; ***P≤0.001). Fold change in mRNA levels were determined as described in the Methods. (B) To examine role of PhoP in *crp* expression, RT-qPCR compared expression levels of *crp* (Rv3676) in WT-H37Rv (empty bar), *ΔphoP*-H37Rv (grey bar) and the complemented mutant (black bar) (***P≤0.001). PhoP-dependent *aprA* expression was shown as a control. (C) *In vivo* recruitment of PhoP within target promoters was examined by ChIP-qPCR as described in the Methods. *espA* promoter (espAup), and *gapdh* - specific enrichments were used as a positive and negative control, respectively. The experiments were performed in biological duplicates, each with two technical repeats (**P≤0.01; ***P≤0.001), and fold enrichment was determined relative to an IP sample without adding antibody (mock control). Non-significant differences are not indicated.

Because the above genes are controlled by *M. tuberculosis* CRP, we wanted to examine if PhoP impacts expression of these genes by controlling *crp* expression. To examine this, expression level of *crp* in WT-, and *ΔphoP*-H37Rv was compared (Fig. 1B). Our results show that *crp* is not under the regulation of *phoP* locus. As a positive control, low pH-inducible *aprA* which belongs to the PhoP regulon, showed a significantly lowered expression in *ΔphoP*-H37Rv relative to WT-H37Rv.

These results suggest that PhoP-dependent regulation of cAMP-inducible/CRP-controlled genes is not attributable to PhoP controlling CRP expression. Next, to investigate whether PhoP is directly recruited within these promoters, Flag-tagged PhoP was ectopically expressed in WT-H37Rv using mycobacterial expression vector p19Kpro [35], and ChIP-qPCR was carried out using anti-Flag antibody as described in the Materials and methods (Fig. 1C). We found that relative to mock sample (no antibody control), PhoP is significantly enriched at the *whiB1*, *icl1* and *umaA* promoters by 12.4±0.4, 4.4±0.5, and 6.2±0.9-fold, respectively. As controls, *gapdh*-specific qPCR did not show any enrichment, whereas *espA* as a previously-reported PhoP regulated promoter [36], showed 6.0±0.3 - fold enrichment relative to the mock sample, further confirming specific recruitment of PhoP within the above CRP-regulated promoters.

### Identifying core binding site of PhoP at the CRP-regulated *whiB1* promoter

Having found PhoP recruitment within these CRP-regulated promoters, we wished to identify the core binding sites of PhoP. To this end, EMSA experiments were carried out using end-labelled whiB1up1 and purified PhoP (Fig. 2). Note that in ChIP assay upstream regulatory region of *whiB1* showed the most efficient *in vivo* recruitment of PhoP (Fig. 1C). In keeping with the ChIP data, P∼PhoP showed ≈20- fold more efficient DNA binding to whiB1up compared to the unphosphorylated regulator (compare lanes 2-4 and lanes 5-7, Fig. 2A [based on limits of detection in these assays]). Additional experiments to identify the core binding site of PhoP using PCR-amplified overlapping promoter fragments show that although 111-bp whiB1up1 (-266 to -156 with respect to the ORF start site of whiB1) efficiently binds to P∼PhoP (lanes 2-3), two other fragments whiB1up2 (-156 to -46), and whiB1up3 (-50 to +60) failed to form a stable complex with P∼PhoP (lanes 5-6, and lanes 8-9, respectively), indicating absence of PhoP binding site downstream of -156 nucleotide of the whiB1 promoter (Fig. S2A). Next, in EMSA experiments, presence of 10- and 20-fold molar excess of unlabelled whiB1up1 efficiently competed out PhoP binding (lanes 3-4, Fig. 2B). However, an identical fold excess of unlabelled whiB1up3 resulted in a minor variation of PhoP binding (<5% effect on DNA binding; lanes 5-6) to whiB1up1 compared to ‘no competitor’ control (lane 2), reflecting that PhoP binding to whiB1up1 is sequence-specific. This result is consistent with the presence of two neighbouring PhoP boxes [22, 25] comprising a 7-bp direct repeat motif spanning nucleotides -97 to -91, and -86 to -80 relative to the *whiB1* transcription start site. To verify the importance of this motif, mutations were introduced within both the repeat units of whiB1up1 as shown in the figure (Fig. 2C). A labelled DNA fragment carrying only these changes (whiB1up1mut) failed to form a stable PhoP - promoter DNA complex, while the WT promoter (whiB1up1) under identical conditions, exhibited efficient DNA binding (compare lanes 2-4 with lanes 6-8, Fig. 2D). These results suggest that PhoP most likely binds to whiB1up1 by sequence-specific recognition of the 7-bp direct repeat motif.

**Fig. 2:**
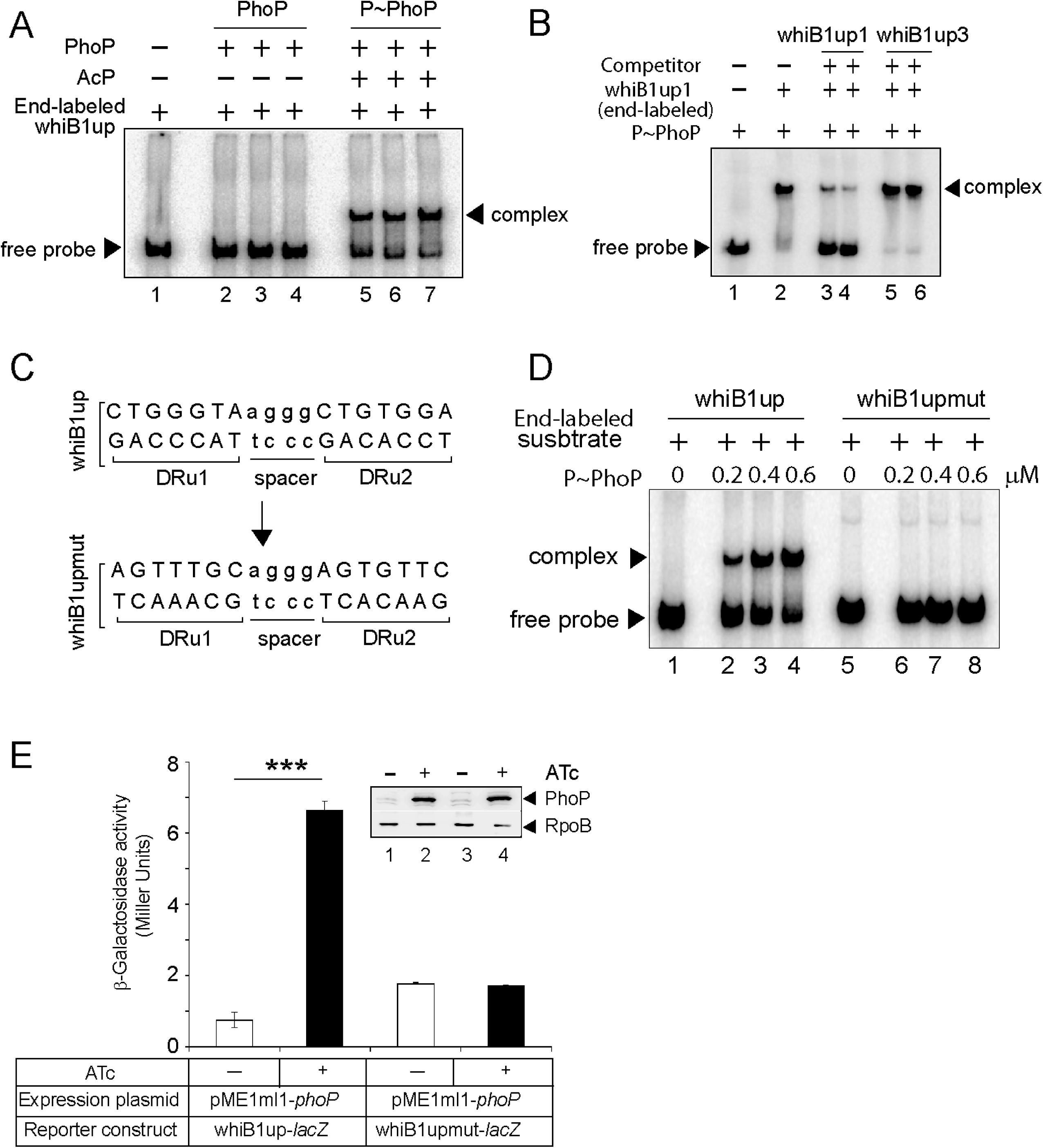
Probing core PhoP binding site within *whiB1* promoter region. (A) EMSA of radio- labelled whiB1up for binding of 0.2, 0.4 and 0.6 µM of PhoP (lane 2-4) or P∼PhoP (lanes 5-7), pre- incubated in a phosphorylation mix with acetyl phosphate (AcP) as the phospho-donor. The arrowheads on the left and right indicate the free probe and a slower moving complex, respectively. To examine sequence-specific binding, EMSA of whiB1up1 with 0.3 µM of P∼PhoP was carried out in absence (lane 2) or in presence of 20, and 40 - fold excess of specific competitor (unlabelled whiB1up1, lanes 3-4) or non-specific competitor (unlabelled whiB1up3, lanes 5-6), respectively. (C) PhoP binding motif consists of upstream (DRu1) and the downstream (DRu2) repeat units. To construct mutant promoter (whiB1upmut), changes in both the repeat units were introduced by changing As to Cs and Gs to Ts and *vice versa*, and the orientation of DRu2 was reversed. whiB1upmut represents whiB1up fragment carrying changes only at the PhoP binding site. (D) EMSA experiment of labelled whiB1up (lanes 2-4), and whiB1upmut (lanes 6-8) to increasing concentrations of P∼PhoP. The free probe and the slower moving complexes are indicated on the figure. The assay conditions, sample analyses, and detection of radio-active samples are described in the Methods. (E) To examine PhoP-regulated expression of whiB1up-*lacZ*, and whiB1upmut-*lacZ* fusions, *M. smegmatis* mc^2^155 harbouring appropriate fusion constructs were grown, and β- galactosidase activities with or without inducing *M. tuberculosis* PhoP expression were measured at 24 hours as described [39]. The results show average values with standard deviations from two biological repeats (\*\*\**P*-value ≤0.001). Inset compares PhoP expression in crude extracts with equal amount of total protein; RpoB was used as a loading control.

To examine the importance of PhoP binding site in transcription regulation, whiB1upmut was cloned into pSM128, a promoter-less mycobacterial reporter plasmid [37] and used as a transcription fusion in *M. smegmatis*. The strain harbouring transcriptional fusions were co-transformed with pME1mL1-*phoP* [38], and grown in 7H9 medium in absence or presence of 50 ng/ml ATc, an inducer of PhoP expression, as described previously [39]. Consistent with *phoP*-dependent *in vivo* activation of *whiB1* (Fig. 1A), with induction of PhoP the whiB1up-*lacZ* fusion was significantly activated by 8.8±0.2-fold at 24 -hour time point as measured by β-galactosidase activity (Fig. 2E). However, under identical conditions, upon induction of PhoP expression cells carrying whiB1upmut*-lacZ* fusion showed a comparable enzyme activity of 0.95±0.03-fold relative to uninduced cells. Differential expression of the regulator in *M smegmatis* impacting promoter activation was ruled out by comparing induction of PhoP expression in bacterial cells carrying either the WT or the mutant promoter (inset to Fig. 2E). Together, we conclude that presence of the newly identified PhoP-binding site, located immediate upstream of CRP binding site, is necessary and sufficient for PhoP-dependent activation of *whiB1*. Based on our existing knowledge on consensus PhoP binding sequence [26, 40], we were able to map likely binding sites of the regulator within the above-noted target promoters (Fig. S2B)

### PhoP promotes CRP recruitment at the *whiB1* promoter

Previous studies had identified CRP binding sites at the *whiB1* promoter [34]. Having discovered PhoP binding site in the close vicinity of CRP binding site, we sought to investigate how a representative promoter DNA accommodates both the regulators. In subsequent binding assays, CRP was unable to bind whiB1up1 to form a complex stable to gel electrophoresis (lanes 2-3, Fig. 3A).

**Fig. 3:**
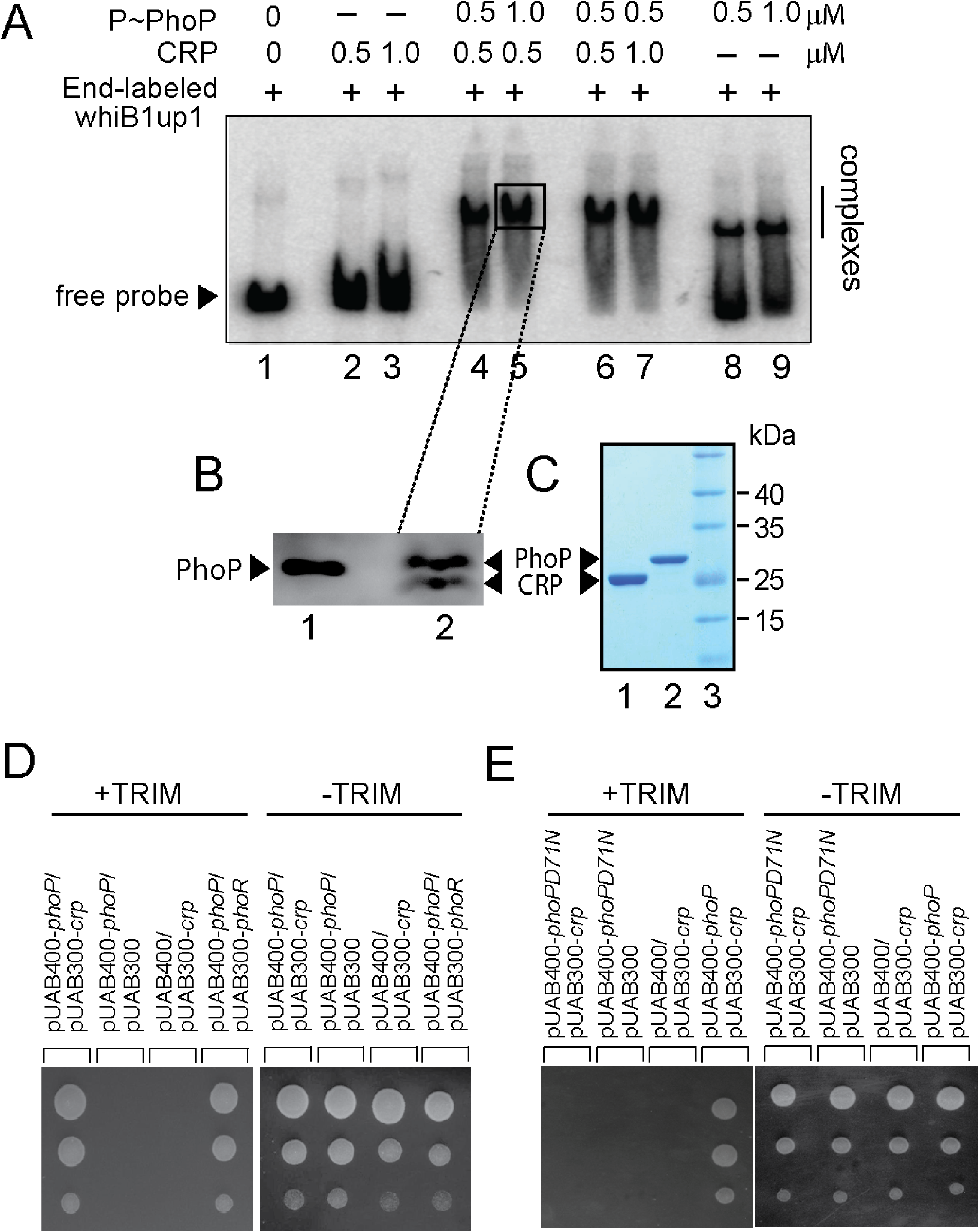
PhoP promotes CRP recruitment at the *whiB1* promoter. (A) To investigate how the promoter simultaneously accommodates both the regulators, EMSA experiments compared end-labelled whiB1up1 binding to increasing concentrations of purified CRP (lanes 2-3), PhoP (lanes 8-9), and both PhoP and CRP together (lanes 4-7). The assay conditions, sample analyses, and detection are described in the Methods section. Positions of the free probe and the complex are indicated on the figure. (B) Western blot analyses of protein fraction extracted from the excised gel fragment representing the complex (lane 5), as indicated by a box, was probed by anti-His antibody in lane 2; lane 1 resolved purified PhoP as a control. (C) Tricine SDS-PAGE analysis was carried out alongside indicating recombinant His-tagged CRP (lane 1) and PhoP (lane 2); lane 3 resolved marker proteins of indicated molecular masses. Protein samples were visualized by Coomassie blue staining. (D-E) To probe protein-protein interaction by mycobacterial protein fragment complementation (M-PFC) assays, *M. smegmatis* expressing either (D) *M. tuberculosis* PhoP and CRP or (E) PhoPD71N (phosphorylation defective PhoP) and CRP, were grown on 7H10/hyg/kan plates in presence of TRIM, and growth was examined for strains co-expressing indicated fusion constructs. In both cases, empty vectors were included as negative controls, and co-expression of pUAB400-*phoP*/pUAB300- *phoR* encoding PhoP and PhoR, respectively, was used as a positive control. All the strains grew well in absence of TRIM.

However, under identical conditions PhoP showed a modest binding to whiB1up1 (lanes 8-9). Interestingly, incubation of both PhoP and CRP together at comparable concentrations with end-labelled whiB1up1, showed a striking stimulation of ≈5-10-fold of DNA binding leading to the formation of a slowest moving band (lanes 4-7). These results demonstrate that the presence of PhoP significantly stimulates CRP recruitment within the *whiB1* promoter. To determine the composition of the ‘slowest moving’ band (lane 5), the protein components of the complex were isolated as described in the Methods, resolved on a Tricine SDS-PAGE and identified by Western blotting using anti His antibody (lane 2, Fig. 3B). Purified regulators were also resolved alongside on a Tricine SDS-PAGE to confirm relative migration of the regulators (Fig. 3C). Our results confirm that the slowest moving band represents a PhoP/CRP/whiB1up1 ternary complex.

Having identified proximal binding sites of CRP and PhoP within *whiB1* promoter, and the two regulators displaying a synergy in DNA binding, we explored the possibility of CRP-PhoP protein-protein interaction using mycobacterial protein fragment complementation (M-PFC) assay (Fig. 3D) [41]. We constructed a bait and prey plasmid pair as C-terminal fusions with complementary fragments of mDHFR. PhoP was expressed from integrative plasmid pUAB400 (Tables S1 and S2 lists oligonucleotide primers and plasmids, respectively). Likewise, *crp* and *phoR* encoding ORFs were expressed from the episomal plasmid pUAB300. The corresponding plasmids were co-transformed in *M. smegmatis* and transformants were selected on 7H10/kan/hyg plates. Interestingly, cells co-expressing PhoP and CRP grew well in presence of TRIM (Fig. 3D). In contrast, *M. smegmatis* harbouring empty vector controls showed no detectable growth on 7H10/TRIM plates while all these strains grew well on 7H10 plates lacking TRIM.

To examine the influence of phosphorylation of PhoP, we next performed M-PFC experiment using phosphorylation-defective PhoPD71N and CRP (Fig. 3E). We found that PhoPD71N is unable to interact with CRP, underscoring the importance of phosphorylation of PhoP on CRP-PhoP interaction. This finding is consistent with phosphorylation impacting DNA binding (Fig. 2A). As a control, under identical conditions, *M. smegmatis* co-expressing PhoP and CMR, cAMP macrophage regulator which regulates a subset of mycobacterial genes in response to variations of cAMP level during macrophage infection [42], did not show any detectable growth (Fig. S3A). To examine the possibility whether variable expression of *M. tuberculosis cmr* and *crp* in *M. smegmatis* contributes to the interaction data, we compared expression levels of the regulators by RT-qPCR (Fig. S3B). Our results suggest that the two regulators CRP and CMR are expressed in *M. smegmatis* at a comparable level, suggesting that altered expression of CMR does not appear to account for lack of growth of the bacteria. Thus, M-PFC data strongly suggest specific interaction between CRP-PhoP.

### Probing CRP-PhoP interactions

To examine the interaction *in vivo*, whole cell lysate of *ΔphoP*-H37Rv expressing His-tagged PhoP was incubated with Ni-NTA as described previously [36]. As expected, both CRP and PhoP were detectable in the crude lysate (lane 1, Fig. 4A). More importantly, upon elution of bound proteins, the eluent showed a clear presence of CRP (lane 3), suggesting specific *in vivo* interaction between CRP and PhoP. As a control, we were unable to detect CRP from the eluent using cell lysates of Δ*phoP*-H37Rv carrying empty vector (p19Kpro) alone (lane 2). Next, *in vitro* pull-down assays were carried out to examine the interaction. In this assay, recombinant PhoP was expressed as a GST fusion protein, and immobilized to glutathione-Sepharose. Following incubation with crude lysate of *E. coli* cells expressing His_6_-tagged CRP, the column-bound proteins were eluted by 20 mM reduced glutathione. Importantly, we could detect presence of both the proteins in the same eluent fraction by immuno-blot analysis (lane 1, Fig. 4B). However, we were unable to detect CRP with only GST- tag (lane 2) or the resin alone (lane 3), allowing us to conclude that PhoP interacts with CRP. We further performed *in vitro* pull-down assays using CRP and phosphorylation-defective PhoPD71N (Fig. S4), as described above. Consistent with the M-PFC data shown in Fig. 3E, our results unambiguously suggest that phosphorylation of PhoP is necessary for CRP-PhoP interaction.

**Fig. 4:**
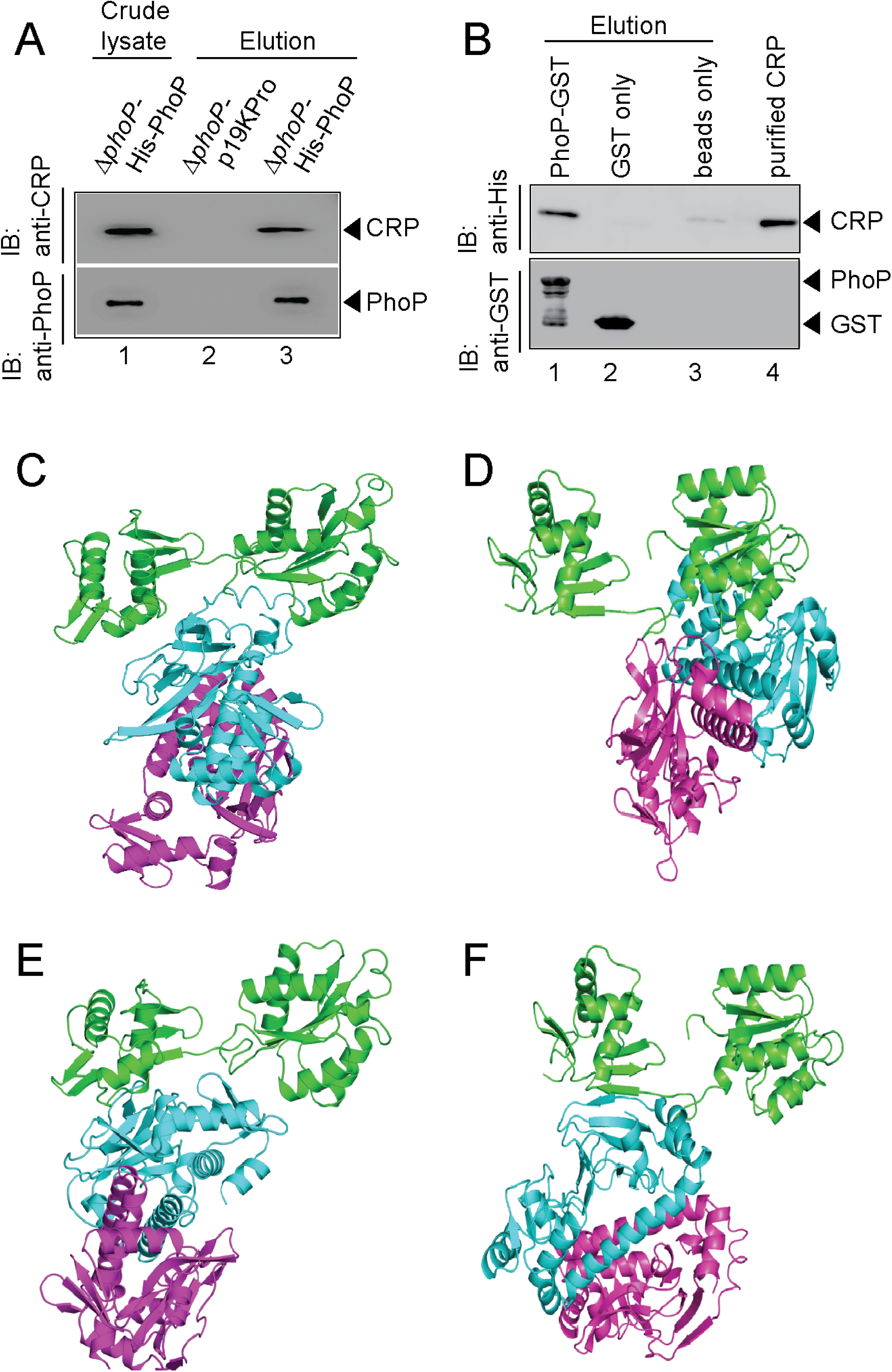
Probing CRP- PhoP interactions. (A) To examine CRP-PhoP interaction *in vivo*, crude cell lysates of *ΔphoP*-H37Rv expressing His_6_-tagged PhoP (p19kpro-*phoP*; Table S2) was incubated with pre-equilibrated Ni-NTA and eluted with 250 mM imidazole; lane 1, input sample; lane 2, control elution from the crude lysate of cells lacking *phoP* expression; lane 3, co-elution of CRP with PhoP. Blots were probed with anti-PhoP and anti-CRP antibody. (B) To investigate CRP-PhoP interaction *in vitro*, crude extract expressing His_6_-tagged CRP was incubated with glutathione sepharose previously immobilized with GST-PhoP. Bound proteins (lane 1) were analyzed by Western blot using anti-His (upper panel) or anti-GST antibody (lower panel). Lane 1 shows presence of CRP bound to GST-PhoP. Identical experiment used glutathione Sepharose immobilized with GST alone (lane 2), or the resin alone (lane 3); lane 4 resolved recombinant His_6_-tagged CRP. (C-F) Rigid body docking to probe CRP-PhoP interaction utilized docking of CRP structure in the free form (PDB ID: 3D0S) or cAMP-bound form (PDB ID: 4A2U) on PhoP (PDB ID: 3 ROJ) by submitting the respective structural coordinates to the docking server ZDOCK as detailed in the ‘Results’ section. Panels A and B show two docked models where a PhoP monomer interacts with a dimeric CRP. Likewise, panels C and D display two docked models where a PhoP monomer interacts with a dimeric cAMP-bound CRP. In these structural models, both PhoP monomers are shown in green; and dimeric CRP, monomers are shown in cyan and magenta, respectively. According to the model of the complex in panel C, PhoPN interacts to the N-domain of CRP. Conversely, panel D suggests involvement of the C-domain of PhoP and CRP with limited involvement of PhoPN. According to panel E, PhoPN interacts with CRP, whereas panel F predicts involvement of the PhoP linker region to CRP-PhoP protein-protein contacts.

To probe structural arrangement of the interacting regulators, X-ray diffracted structures of PhoP (PBDID: 3ROJ) and a dimer of CRP were docked employing ZDOCK [43] (Figs. 4C-F). *M. tuberculosis* CRP structure is available both in free- (PDB ID: 3D0S) and cAMP-bound form (PDB ID: 4A2U). Although these structures are nearly identical (RMSD less than 0.1A° for backbone atoms), cAMP induces a conformational change in the C-terminal DNA binding domain [44]. Therefore, CRP-PhoP rigid-body-docking utilized structural coordinates of both free (Fig. 4C-D) and cAMP-bound CRP (Fig. 4E-F). The missing amino acids of PhoP structure (residues 142-147) were modeled using MODELLER [45, 46], the docked complexes were ranked using ZRANK [47], and top-ranked complexes were identified using PDBePISA webserver [48]. Next, nearly two thousand top scoring complexes were screened, and further re-ranked using a scoring function to yield six docked complexes. Two possible models of interaction emerged based on the above analyses. The first one of the two (based on free CRP) suggests interaction between the dimerization domain of PhoP with CRP (Fig. 4C), whereas the second model predicts that the C-domain of PhoP contacts CRP (Fig. 4D). Conversely, using cAMP-bound CRP as the structural template, ZDOCK predicted two possible interaction models. While first one of the two suggests involvement of the N-domain of PhoP (Fig. 4E), the other model is indicative of involvement of the PhoP linker in contacting CRP (Fig. 4F). In summary, docking analysis suggests that it is the N-terminal domain of CRP, which appears to interact with PhoP. In contrast, away from the interaction interface, the C-terminal domain remains available to interact with DNA, arguing in favour of the fact that absence of cAMP is unlikely to influence interaction interface of CRP. These observations are consistent with previous results suggesting a significantly lesser effect of cAMP on functioning of *M. tuberculosis* CRP relative to its *E. coli* orthologue [49], and provide a justification why cAMP was not included in our experiments investigating CRP-PhoP interaction.

### CRP and PhoP interact via their corresponding N-terminal domains

To examine between the two possibilities, we assessed role of different stretches of PhoPN in CRP-PhoP interactions (Fig. 5A). *In vitro* pull-down assays using mutant GST-PhoP proteins (each with three potential CRP- contacting residues of PhoP replaced with Ala) with His-tagged CRP displayed effective protein-protein interaction as that of the WT PhoP (compare lane 1 and lanes 2-6). These results suggest that either PhoPN does not contribute to CRP-PhoP interactions or possibly a constellation of residues, but not a few residues of PhoPN, in a single stretch is involved in CRP-PhoP interaction(s). Next, pull-down assays using His-tagged CRP and a linker deletion mutant of PhoP [PhoPLAla5, replacing 5 residues spanning Gly^142^ to Pro^146^ with Ala, which remains functional [50], suggest that PhoP linker does not appear to contribute to CRP-PhoP interaction(s) (compare lane 1 and lane 2, Fig. 5B). We next probed CRP-PhoP interaction using His-tagged full-length CRP and PhoP domains as GST-fusion constructs (Fig. 5C). PhoPN and PhoPC were previously shown to be functional for phosphorylation and DNA binding activity, respectively [50] on their own. Importantly, PhoPN (comprising residues 1-141) showed effective protein-protein interaction with CRP (lane 1). However, PhoPC (comprising residues 141-247), under identical conditions, did not display an effective interaction with CRP (lane 2). To identify the corresponding interacting domain of CRP, we next used GST-PhoP and His-tagged CRP-domain constructs (Fig. 5D). Interestingly, N-terminal domain of CRP, CRPN comprising residues 28-116 co-eluted with GST-PhoP (lane 1). However, C-terminal domain of CRP, CRPC comprising residues 146-224 under identical conditions, did not co-elute with GST-PhoP (lane 2), suggesting specific interaction between CRPN and PhoP. Together, we surmise that CRP-PhoP interaction is mediated by the N-domains of the corresponding regulators.

**Fig. 5:**
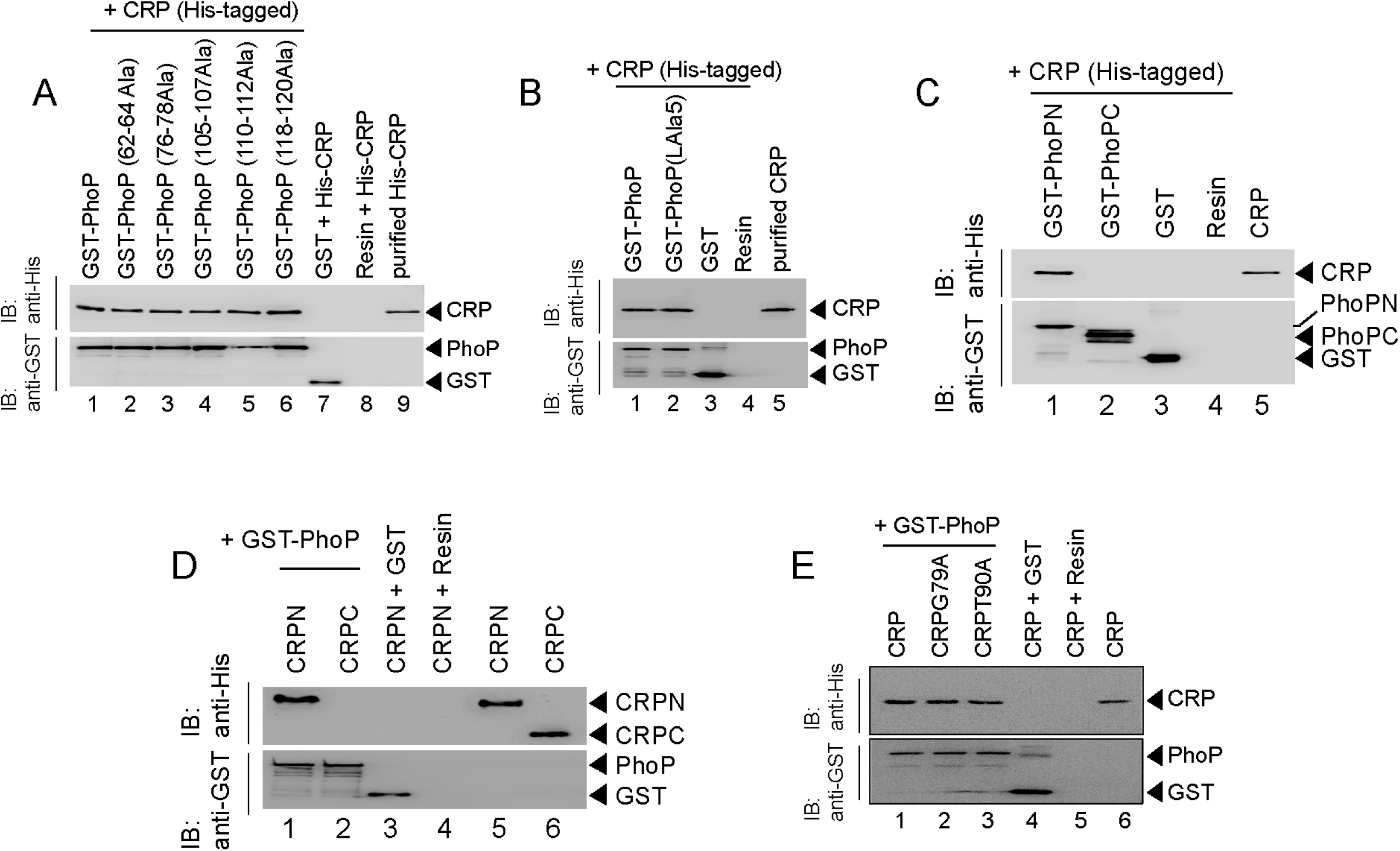
PhoP and CRP interact with each other via their corresponding N-terminal domains. (A) To assess the importance of residues of PhoPN, crude lysates of cells expressing His_6_-tagged CRP was incubated with glutathione-Sepharose, previously immobilized with GST-tagged WT (lane 1) or mutant PhoP proteins (lanes 2-6), each carrying a substitution of three PhoP residues with alanine (as indicated on the figure). Fractions of bound proteins (lane 1-6) were analysed by Western blot using anti-His (upper panel) or anti-GST antibody (lower panel). Control sets include glutathione Sepharose immobilized with GST alone (lane 7), or the resin alone (lane 8); lane 9 resolved recombinant His_6_- tagged CRP. (B) To examine role of the PhoP linker in CRP-PhoP interactions, crude lysates of cells expressing His_6_-tagged CRP was incubated with glutathione-Sepharose previously immobilized with GST-tagged PhoP (lane 1) or a PhoP linker mutant (lane 2), carrying a substitution of five linker residues (spanning Gly^142^ to Pro^146^ of PhoP) with alanine as described previously [50]. Analysis of bound fractions (lanes 1-2) was carried out as above and control sets include glutathione Sepharose immobilized with GST alone (lane 3), or the resin alone (lane 4); lane 5 resolved recombinant CRP. (B) To examine importance of cAMP binding to CRP on CRP-PhoP interaction, *in vitro* pull-down assays were carried out using His-tagged CRP mutants, deficient for cAMP binding. The experimental procedures, and data analyses are as described in the legend to Fig. 4B. (D-E) CRP-PhoP interaction was probed by *in vitro* pull-down assays using either (D) His-tagged CRP and GST-tagged PhoP domains (GST-PhoPN and GST-PhoPC, respectively) or (E) GST-tagged PhoP and His-tagged CRP domains (His-CRPN, and His-CRPC, respectively). The domain constructs are listed in Table S2. Fractions of bound proteins were analysed by Western blot using anti-His (upper panel) or anti-GST antibody (lower panel). Control sets include glutathione Sepharose immobilized with GST (lane 3), or the resin (lane 4) alone; lane 5 of panel A resolved purified CRP, while lanes 5, and 6 of panel B resolved purified CRPN, and CRPC, respectively.

Although CRP-PhoP interaction studies were performed in the absence of cAMP, we next investigated whether cAMP binding to CRP is required for this protein-protein interaction. Thus, we performed *in vitro* interaction studies using CRP mutants (CRPG79A, and CRPT90A), deficient for cAMP binding (Fig. 5E). The mutations were designed based on previously-reported CRP structure defining the cAMP binding pocket of the regulator [51]. Our pull-down assays demonstrate that the mutant CRP proteins interact with PhoP as effectively as that of the wild-type CRP, allowing us to conclude that cAMP binding to CRP is not required for CRP-PhoP interaction. Importantly, these observations are consistent with docking studies involving both free and cAMP-bound CRP structures (Fig. 4). Further, in keeping with cAMP-independent functioning of *M. tuberculosis* CRP [6, 49, 52], the mutant proteins over a range of concentrations showed effective DNA binding to end-labelled whiB1up with a comparable affinity as that of the wild-type CRP (Fig. S5). These results also suggest that the mutant proteins retain native structure and/or fold as that of wild-type CRP.

### Functioning of CRP requires presence of PhoP

The CRP regulon was previously studied by comparing transcription profiling of WT-H37Rv and a CRP-depleted strain of *M. tuberculosis* H37Rv *(Δcrp*-H37Rv) using RNA-seq and ChIP-seq [21]. To examine the influence of PhoP, we next undertook to investigate *in vivo* recruitment of CRP within CRP-regulated promoters of WT-and *ΔphoP*-H37Rv. Formaldehyde-cross-linked DNA-protein complexes of growing *M. tuberculosis* cells were sheared to generate fragments of average size ≍500 bp as described previously [53]. In ChIP experiments using anti-CRP antibody, DNA binding was analysed by qPCR (Fig. 6A). Our results show that relative to mock sample, CRP is effectively recruited within its target promoters. For example, *icl1*, and *whiB1* promoters, displayed 5.6±1 and 16.4±2.8**-**fold enrichments of qPCR signals, respectively. In contrast, under identical conditions *ΔphoP*-H37Rv -derived samples showed insignificant enrichments of CRP within *icl1* (0.7±0.2-fold) and *whiB1* (0.6±0.18-fold), promoters, respectively. However, with identical input DNA sample, we observed a largely comparable CRP recruitment within the CRP-regulated *sucC* promoter (sucCup, which does not belong to PhoP regulon) [21] in both the WT- and the mutant bacilli. Inset shows a comparable expression of CRP in WT-H37Rv and *ΔphoP*-H37Rv. These results strongly suggest that both mycobacterial regulators are recruited concomitantly within a subset of CRP-regulated promoters and consistent with CRP-PhoP interaction, absence of PhoP strikingly influences *in vivo* recruitment of CRP.

**Fig. 6:**
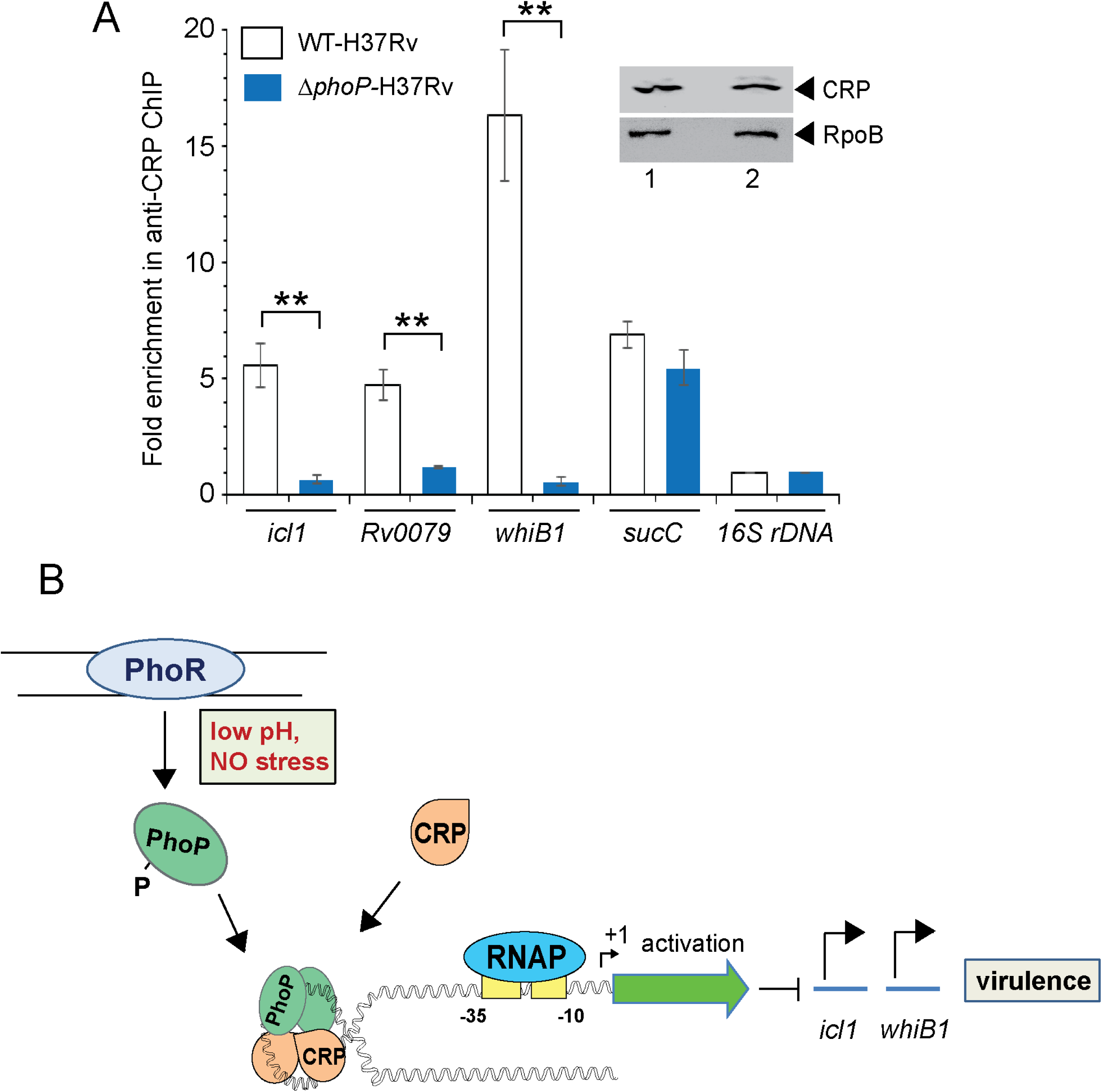
PhoP promotes CRP recruitment to regulate *whiB1* expression. To examine effect of CRP- PhoP interaction, chromatin-immunoprecipitation (ChIP)-qPCR was carried out using anti-CRP antibody to compare *in vivo* recruitment of CRP in WT-H37Rv and *ΔphoP*-H37Rv as described in the Methods. Fold PCR enrichment due to CRP binding to indicated promoters was determined by duplicate measurements, each with a technical repeat (**P≤0.01). Inset compares CRP expression in ≍10 μg of indicated crude cell-lysates as probed by anti-CRP antibody; identical extracts were probed with anti-RpoB antibody as a loading control. (B) Schematic model showing newly-proposed molecular mechanism of activation of CRP-regulated promoters by simultaneous binding of CRP and PhoP. It is noteworthy that CRP has been suggested to function as a general chromosomal organizer [11]. We propose that the additional stability of the transcription initiation complex is perhaps derived from the protein-protein interaction between CRP and PhoP. While these interacting proteins remain bound to their cognate sites away from the start site(s), and contribute to the stability of the transcription initiation complex, RNA polymerase (RNAP) effectively transcribes these genes. Together, these molecular events mitigate stress by controlling expression of numerous genes and perhaps contribute to better survival of the bacilli in cellular and animal models..

## Discussion

*M. tuberculosis*, as a successful intracellular pathogen heavily relies on its ability to sense and appropriately respond to varying environmental conditions over the course of an infection. cAMP, one of the most widely used second messengers, impacts on a wide range of cellular responses in mycobacterial physiology including host-pathogen interactions and virulence [8]. In addition, a large number of genome-encoded adenylate cyclases in *M. tuberculosis* complex is remarkable in view of a single adenylate cyclase of most bacteria. Consistent with the abundance of adenylate cyclases, mycobacteria produce high levels of cAMP relative to other bacterial species [54]. Importantly, cAMP levels are elevated upon infection of macrophages by pathogenic mycobacterium [54] and addition of cAMP to growing cultures of *M. tuberculosis* significantly impacts mycobacterial gene expression [4]. Agarwal and colleagues had shown that bacterial cAMP burst, upon infection of macrophages, improves bacterial survival by interfering with host signalling pathways [9]. However, mechanism of activation of cAMP-responsive/CRP-regulated mycobacterial genes remain poorly understood. This study identified at least part of CRP regulon in *M. tuberculosis*, and established PhoP in addition to CRP and CMR as a third cAMP-responsive transcription factor. Notably, PhoP was required for the regulated expression of a subset of CRP-regulated genes. The CRP-PhoP associated regulon is distinct from the reported CRP regulon [21, 55] with respect to their mechanism of transcription activation, and most likely responds to environmental conditions that regulate its expression.

Having noted that numerous genes regulated by CRP also belong to PhoP regulon and the *whiB1* promoter region displays canonical CRP and PhoP binding sites, we sought to explore a functional link between CRP and PhoP. Importantly, presence of PhoP binding site (Fig. 2C) was consistent with *in vivo* recruitment of PhoP within CRP-regulated promoters (Fig. 1C). Using *in vitro* and *in vivo* approaches, we showed that CRP interacts with PhoP (Figs. 3-4). While phosphorylation of PhoP was found to be necessary for CRP-PhoP interaction (Fig. 3E), consistent with cAMP-independent functioning of CRP [6, 49, 52], mutant CRP proteins defective for cAMP binding showed effective CRP-PhoP interaction (Fig. 5C). To examine how a representative promoter recruits both the regulators *in vitro*, we attempted an EMSA experiment using purified regulators (Fig. 3A).

The fact that CRP and PhoP together form a ternary complex at the *whiB1* promoter, prompted us to investigate a possible inter-dependence of recruitment of the two regulators. Despite effective recruitment of CRP in WT bacilli, strikingly CRP recruitment within identical promoters was almost undetectable in *ΔphoP*-H37Rv (Fig. 6A), a result which finds an explanation from CRP-PhoP interaction data (Figs. 3-4). Given the importance of CRP-regulated essential genes like *whiB1*, and *icl1* in virulence, CRP-PhoP interactions controlling *in vivo* regulation of expression provides one of the most fundamental biological insights into the mechanism of virulence regulation of *M. tuberculosis*. The finding that PhoP functions as a prerequisite to ensure specific recruitment of CRP within a few CRP-regulated promoters, offers a new biological insight into the controlled expression of CRP regulon. Because such a situation might assist PhoP to function efficiently by ensuring CRP binding only to target promoters already bound to PhoP, it is conceivable that PhoP binding changes DNA conformation, which facilitates CRP recruitment. Therefore, we propose that a DNA-dependent complex interplay of CRP and PhoP via protein-protein contacts most likely controls precise regulation of these genes. Indeed, this model fits well with, explains and integrates previously reported regulations of *icl1* and *whiB1*, independently by PhoP [23, 30] and CRP [21], respectively.

Based on our observations that the N-domains of PhoP and CRP interact to each other (Figs. 5C-D), and effective CRP recruitment is possibly ensured within target promoters already bound to PhoP (Fig. 6A), our model suggests that CRP-PhoP interaction(s) account for activation of CRP-regulated genes. Although we do not provide evidence to suggest 1:1 binding stoichiometry except that as independent regulators, in both cases a dimer of PhoP [31, 32] and CRP [44] bind to target DNA, these considerations take on more significance in the light of previously-identified CRP binding site [6, 49] and newly-identified PhoP binding site (this study) within the *whiB1* promoter.

The arrangement and spacing between the two binding sites are suggestive of DNA looping, possibly to assist transcription initiation by contributing additional stability to the initiation complex. Having noted *in vivo* relevance of CRP-PhoP interaction impacting a few promoters, we sought to explore whether this is a more generalized mechanism involving numerous other CRP-regulated mycobacterial genes. A careful inspection of ChIP-seq data identifying genome- wide CRP binding sites [21] and SELEX and ChIP-seq data of PhoP [22, 25] remarkably uncover presence of ∼10 such promoters comprising CRP and PhoP binding sites (Table 1). These results suggest a critically important role of PhoP in binding and transcriptional control of CRP-regulon genes and establish a molecular mechanism showing how two regulators function as co-activators of gene expression in response to phagosomal stress. Given the complex life cycle of the intracellular pathogen which encounters a variety of physiological conditions within the human host, an integration of more than one regulator might be required for *in vivo* functioning to achieve a rather complex regulation of these mycobacterial genes. While this may be possibly a more common strategy than has been previously appreciated, in *M. tuberculosis* essential gene like *whib1* [34], virulence determinant like *icl1* [56–58], which remain critical for a successful infection, most likely are under precise regulation as deregulated expression of these genes is likely to alert the immune system to the infection.

**Table 1.**
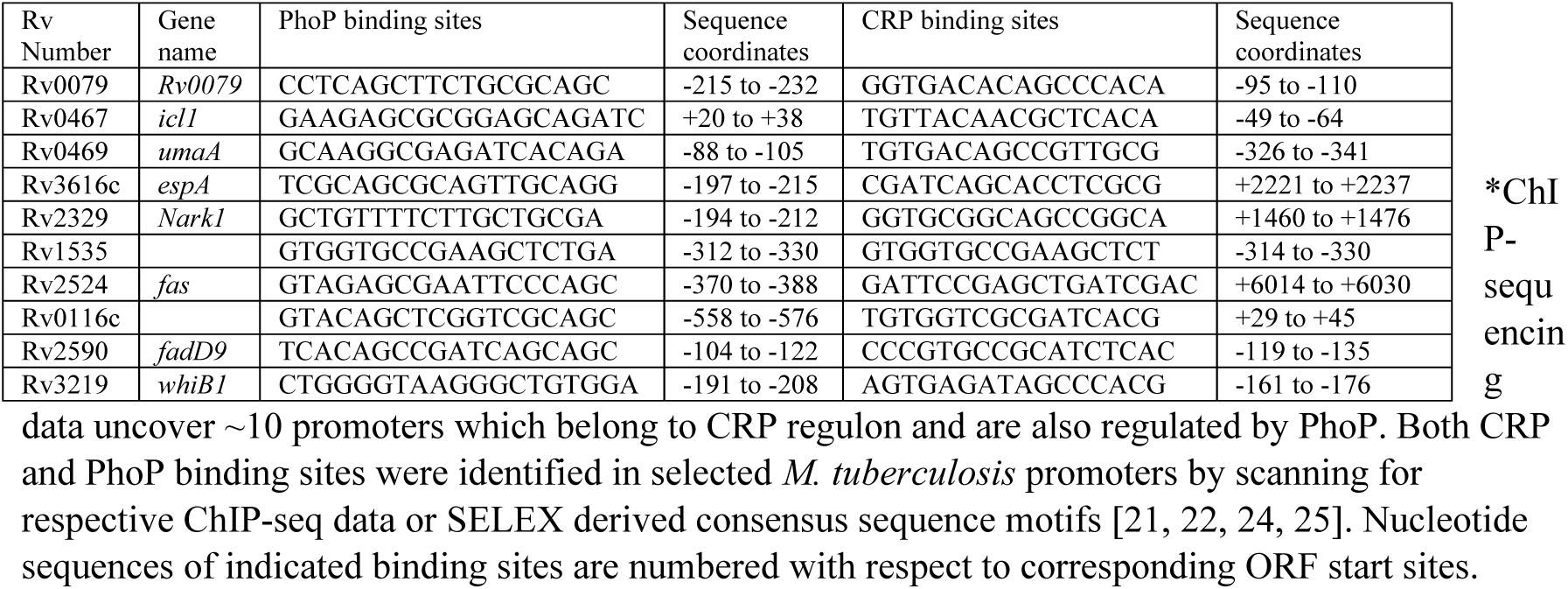
CRP and PhoP binding sites within the commonly regulated promoters*

In conclusion, the present study identifies a new mechanism and provided critically new insights into CRP-dependent gene regulation by the two interacting virulence factors. Future work is required to better understand the components of signalling pathways that makes use of this particular mechanism of gene regulation. In addition, the broader questions of whether CRP-dependent mycobacterial gene regulation is triggered by cAMP produced from a specific adenyl cyclase, and whether in response to specific environmental changes, are likely to be exciting areas of future investigation.

## Materials and Methods

### Bacterial strains and culture conditions

The strains used were *E. coli* DH5α, for all cloning experiments; *E. coli* BL21 DE3, for protein expression; *M. smegmatis* mc^2^155, for mycobacterial- protein-fragment complementation (M-PFC), and reporter-based transcription regulation experiments; wild-type *M. tuberculosis* H37Rv (WT-H37Rv) and *ΔphoP*-H37Rv, in which *phoPR* locus (Rv0757- Rv0758) has been inactivated [23], for the results reported in this study. *M. tuberculosis* H37Rv, its derivatives and *M. smegmatis* mc^2^155, were grown at 37°C in Middlebrook 7H9 liquid broth (Difco) containing 0.2% glycerol, 0.05% Tween-80 and 10% ADC (albumin-dextrose-catalase) or on 7H10- agar medium (Difco) containing 0.5% glycerol and 10% OADC (oleic acid-albumin-dextrose- catalase). Growth, transformation of mycobacterial strains and selection of transformants on appropriate antibiotics were carried out as described [39]. For *in vitro* growth under specific stress conditions, indicated mycobacterial strains were grown to mid log phase (OD_600_ 0.4-0.6) and exposed to different stress conditions. For acid stress, cells were initially grown in 7H9 media, pH7.0 and on attaining mid log phase it was transferred to acidic media (7H9 media, pH 4.5) for further two hours at 37°C. For oxidative stress, cells were grown in presence of 50 µM CHP (Sigma) for 24 hours or indicated diamide concentration(s) for 7 days. For NO stress, cells grown to mid log phase were exposed to 0.5 mM DataNonoate for 40 minutes [59].

### Cloning

*M. tuberculosis* full-length *crp* (Rv3676; encoded by 672-bp of the ORF), truncated N- terminal domain *crpN* (encoded by 267-bp of the ORF), and *crpC* (encoded by 237-bp of the ORF) over-expressing constructs were cloned in T7-lac-based expression system pET28c (Novagen) between NdeI and HindIII sites as recombinant fusion proteins (containing an N-terminal His_6_-tag) using primer pairs FPcrp/RPcrp, FPcrpN/RPcrpN, and FPcrpC/RPcrp (Table S1) resulting in plasmids as listed in Table S2. Plasmid pGEX-*phoP* expressed GST-PhoP, the full-length PhoP protein with an N-terminal GST-tag [32]. Likewise, pGEX-*phoPN* and pGEX-*phoPC,* generated by cloning of corresponding ORFs between BamHI and XhoI sites of pGEX 4T-1 (GE Healthcare), expressed PhoPN, and PhoPC, respectively, each with an N-terminal GST tag. To express *Rv0805*, the ORF was cloned between EcoRV and HindIII sites of the mycobacterial expression vector pSTKi [60] and expressed from the P_myc1_*tetO* promoter. Mutations in CRP was introduced by two-stage overlap extension method using mutagenic primers (Table S1), and each construct was verified by DNA sequencing.

### Proteins

Full-length and truncated PhoP proteins containing either an N-terminal His_6_- tag or an N- terminal GST-tag, were purified as described previously [32]. Wild-type CRP protein (Rv3676) from *M. tuberculosis* H37Rv was expressed in *E. coli* BL21(DE3) as a fusion protein containing an N- terminal His_6_-tag (Novagen) and purified by metal-affinity chromatography (Ni-NTA, Qiagen). The protein expression was induced by adding 0.4 mM IPTG in log-phase cultures at O.D_600_ of 0.4, and cells were allowed to grow overnight at 16°C. All subsequent procedures were carried out at 4°C. Briefly, cells were resuspended in lysis buffer (50 mM Tris-HCl, pH 7.9, 500 mM NaCl, 10% glycerol, 0.25% Tween-20, and 50 mM imidazole) followed by addition of lysozyme to a final concentration of 0.1 mg/ml. Next, cell lysates were sonicated, and insoluble material was removed by centrifugation for 30 min at 12,000 rpm. The clear supernatant was applied to a column of Ni-NTA agarose (Qiagen) that had been equilibrated with the lysis buffer. After an incubation of 30 minutes, the column was repetitively washed with 10 column volumes of lysis buffer only, and then with 1 column volume of each of lysis buffer containing 1M NaCl, lysis buffer containing 50 mM imidazole, and lysis buffer containing 100 mM imidazole, respectively. Finally, the bound protein was eluted in lysis buffer containing 600 mM imidazole. Before storage at -80°C, the protein was extensively dialyzed against buffer A (50 mM Tris-HCl, pH 7.9, 300 mM NaCl, and 10% glycerol), protein concentration was determined using Bradford reagent with BSA as the standard, and expressed in equivalent of protein monomers.

### RNA isolation

Total RNA was extracted from exponentially growing bacterial cultures grown with or without specific stress as described above. Briefly, 25 ml of bacterial culture was grown to mid-log phase (OD_600_= 0.4 to 0.6) and combined with 40 ml of 5 M guanidinium thiocyanate solution containing 1% β-mercaptoethanol and 0.5% Tween 80. Cells were pelleted by centrifugation, and lysed by re-suspending in 1 ml Trizol (Ambion) in the presence of Lysing Matrix B (100 µm silica beads; MP Bio) using a FastPrep-24 bead beater (MP Bio) at a speed setting of 6.0 for 30 seconds. The procedure was repeated for 2-3 cycles with incubation on ice in between pulses. Next, cell lysates were centrifuged at 13000 rpm for 10 minutes; supernatant was collected and processed for RNA isolation using Direct-Zol^TM^ RNA isolation kit (ZYMO). Following extraction, RNA was treated with DNAse I (Promega) to degrade contaminating DNA, and integrity was assessed using a Nanodrop (ND-1000, Spectrophotometer). RNA samples were further checked for intactness of 23S and 16S rRNA using formaldehyde-agarose gel electrophoresis, and Qubit fluorometer (Invitrogen).

### Quantitative Real-Time PCR

cDNA synthesis and PCR reactions were carried out using total RNA extracted from each bacterial culture, and Superscript III platinum-SYBR green one-step qRT-PCR kit (Invitrogen) with appropriate primer pairs (2 µM) using an ABI real-time PCR detection system. Oligonucleotide primer sequences used in RT-qPCR experiments are listed in Table S3. Control reactions with platinum Taq DNA polymerase (Invitrogen) confirmed absence of genomic DNA in all our RNA preparations, and endogenously expressed *M. tuberculosis rpoB* was used as an internal control. Fold difference in gene expression was calculated using ΔΔC_T_ method [61]. To determine enrichment due to PhoP and/or CRP binding targets in the IP DNA samples, 1 µl of IP or mock IP (no antibody control) DNA was used with SYBR green (Invitrogen) along with target promoter-specific primers. Average fold differences in mRNA levels were determined from at least two biological repeats each with two technical repeats. Non-significant difference is not indicated.

### ChIP-qPCR

ChIP experiments were carried out with some modifications of the protocol described previously [62]. To determine PhoP recruitment, FLAG tagged *phoP* ORF was expressed in WT- H37Rv and *ΔphoP*-H37Rv from mycobacterial expression vector p19Kpro [35], and ChIP was carried out by anti-FLAG antibody. To determine CRP recruitment by ChIP, the CRP specific antibody was used. *M. tuberculosis* CRP (Rv3676), tagged with His_6_ at the N-terminus was purified in *E. coli* and used to produce CRP-specific polyclonal antibody in rabbit by AlphaOmegaSciences (India). 0.3 mg of purified recombinant protein emulsified in Freund’s complete adjuvant was administered as primary dose subcutaneously, followed by two booster immunizations in Freund’s incomplete adjuvant after 14 and 30 days into New Zealand White Rabbit (∼ 3.8 kg body weight). The titre was determined by the endpoint method, blood collected, serum separated and stored at -20°C. Total Immunoglobulin G (IgG) was purified from the serum by affinity chromatography using Protein-A resin].

For ChIP assays mycobacterial cells were grown to mid exponential phase (OD_600_≍0.4-0.6) and formaldehyde was added to a final concentration of 1%. After incubation of 20 minutes, glycine was added to a final concentration of 0.5 M to quench the reaction and incubated for further 10 minutes. Cross-linked cells were harvested by centrifugation and washed twice with ice-cold immunoprecipitation (IP) buffer [50 mM Tris (pH 7.5), 150 mM NaCl, 1 mM EDTA, 1% Triton X- 100, 1 mM PMSF and 5% glycerol]. Cell pellets were resuspended in 1.0 ml IP buffer containing protease inhibitor cocktail (Roche), cells were lysed and insoluble matter was removed by centrifugation at 13000 rpm for 10 min at 4°C. Next, supernatant, containing DNA, was sheared to an average size of ∼500 base pairs (bp) using a Bioruptor (Sonics, VibraCell) with settings of 20s on and 40s off for 3-5 minutes, and split into two aliquots. 10µl of sample was analysed on agarose gel to check the size of the DNA fragments. Each ∼0.5 ml supernatant was incubated with either no antibody (mock-IP), or 100 µg specific antibody at 4°C overnight. In a parallel set up, 25 µl Protein A beads were incubated with 50 µg herring sperm DNA in 500 µl of IP buffer on a rotary shaker at 4°C overnight. Next, samples with or without antibody were individually added to the tubes containing Protein A beads and incubated at 4°C overnight. Finally, samples were washed twice with IP buffer, twice with IP buffer containing 500 mM NaCl, once with wash buffer [10 mM Tris (pH 8.0), 250 mM LiCl, 1 mM EDTA, 0.5% Tergitol (Sigma) and 0.5% sodium deoxycholate] and once with TE (pH 7.5). IP complexes were then eluted from the resin in 350 µl elution buffer [50 mM Tris (pH 7.5), 10 mM EDTA, 1% SDS and 100 mM NaHCO_3_] containing 0.8 mg*/*ml Proteinase K at 65°C for overnight. The resulting IP samples were then ethanol precipitated.

*In vivo* recruitment of the regulators were determined using appropriate dilutions of IP DNA in a reaction buffer containing SYBR green mix (Invitrogen), 2 µM PAGE-purified primers (Table S3) and one unit of Platinum Taq DNA polymerase (Invitrogen). Typically, 40 cycles of amplification were carried out using real time PCR detection system (Eppendorf). qPCR signal from an IP experiment without adding an antibody (mock) was measured to determine the efficiency of recruitment. In all cases, melting curve analysis confirmed amplification of a single product. Specificity of PCR-enrichment from the identical IP samples was verified using 16S rDNA- specific primers. Each data was collected in duplicate qPCR measurements using at least two bacterial cultures.

### EMSA

The DNA probes were generated by PCR amplification of whiB1up, resolved on agarose gels, recovered by gel extraction, end-labelled with [γ-^32^P ATP] (1000 Ci nmol^-1^) using T4 polynucleotide kinase and purified from free label by Sephadex G-50 spin columns (GE Healthcare). Increasing amounts of purified regulators were incubated with appropriately end-labelled DNA probes in a total volume of 10 µl binding mix (50 mM Tris-HCl, pH 7.5, 50 mM NaCl, 0.2 mg/ml of bovine serum albumin, 10% glycerol, 1 mM dithiothreitol, ≈ 50 ng of labeled DNA probe, and 0.2 µg of sheared herring sperm DNA) at 20°C for 20 minutes. DNA-protein complexes were resolved by electrophoresis on a 6% (w/v) polyacrylamide gel (non-denaturing) in 0.5X TBE (89 mM Tris-base, 89 mM boric acid and 2 mM EDTA) at 70 V and 4°C, and radioactive bands were quantified by the phosphorimager (Fuji). To identify composition of retarded complexes, the position of the radioactive material was determined by exposure to a phosphor storage screen, and bands representing the complex(es) were excised from the gel. Next, protein components were extracted from the excised gel fragments in buffer contaning 50 mM Tris, pH 7.5, 50 mM NaCl, 10% glycerol and 1 mM DTT as described previously[36]. The eluted protein samples were concentrated, resolved in tricine SDS-PAGE, and detected by Western blotting using appropriate anitbodies.

### Mycobaterial protein fragment complementation (M-PFC) assay

*M. tuberculosis* PhoP was cloned in pUAB400 (kan^R^; Table S2) and pUAB400-*phoP* expressed in *M. smegmatis* as described previously [53]. Transformed cells were selected on 7H10/ kan plates and grown in liquid medium to obtain competent cells of *M. smegmatis* harbouring pUAB400-*phoP*. Likewise, *crp* and *cmr* encoding genes were amplified from *M. tuberculosis* H37Rv genomic DNA using primer pairs FPmCRP/RPmCRP, and FPmCMR/RPmCMR, respectively (Table S1), and cloned in episomal plasmid pUAB300 (hyg^R^; Table S2) between PstI/HindIII and BamHI/HindIII sites, respectively. Each construct was verified by DNA sequencing. Next, co-transformants of *M. smegmatis* were selected on 7H10/kan/hyg plates both in absence and presence of 10 µg/ml of TRIM to investigate protein-protein interactions [53]. As a positive control, plasmid pair expressing *phoP*/*phoR* was used as described previously [62].

## Acknowledgements

We thank G. Marcela Rodriguez and Issar Smith (PHRI, New Jersey Medical School - UMDNJ) for Δ*phoP*-H37Rv, and the complemented *M. tuberculosis* H37Rv strains, Adrie Steyn (University of Alabama) for pUAB300/pUAB400 plasmids, and Ashwani Kumar (CSIR-IMTECH) for helpful discussions. We acknowledge Pragya Priyadarshini for her assistance in docking studies, and Sanjeev Khosla for critical reading of the manuscript. This study received financial support from intramural grants of CSIR (MLP-0049) and CSIR-IMTECH (OLP-0170), and by a research grant (to D.S) from SERB (EMR/2016/004904), Department of Science and Technology (DST). H.K., P.P., R.R.S., and S.K were supported by CSIR pre-doctoral fellowships.

## Author contributions

H.K., P.P., R.R.S., and D.S. designed research; H.K., P. P., R.R.S., and S. K. performed research; B.S. contributed analytical tools; H.K., P. P., R.R.S., B.S. and D.S. analysed data; and D.S. wrote the paper.

## Supplemental Data

The supplemental data include five supplemental figures (Figs. S1-S5), and three supplemental tables (Tables S1-S3). All other relevant data are part of the main text and its supplemental files.

## Competing interests

The authors declare that no competing interests exist.

## Supplemental Information

**Table S1.**
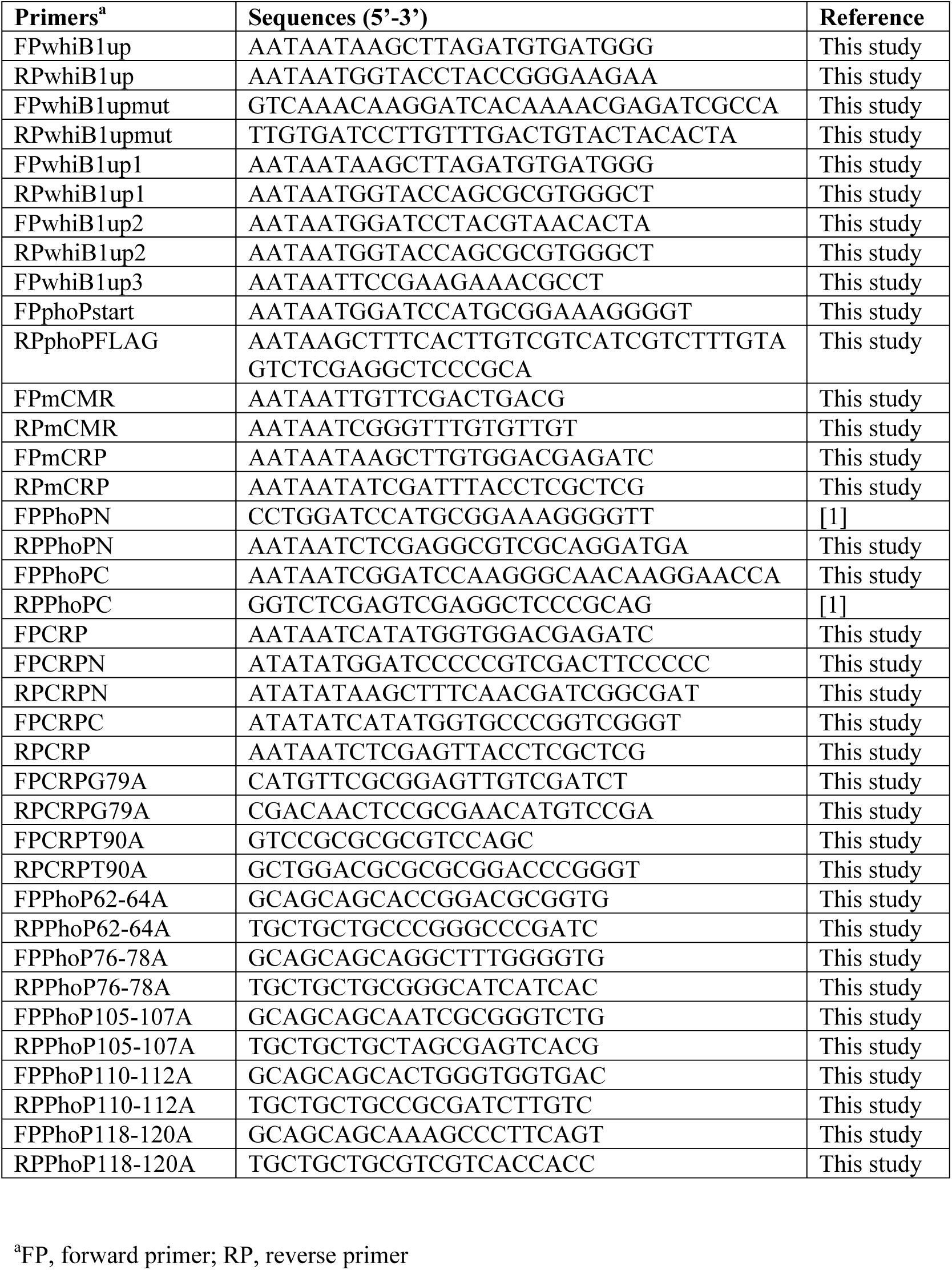
Oligonucleotide primers used for amplification and cloning in this study

**Table S2.**
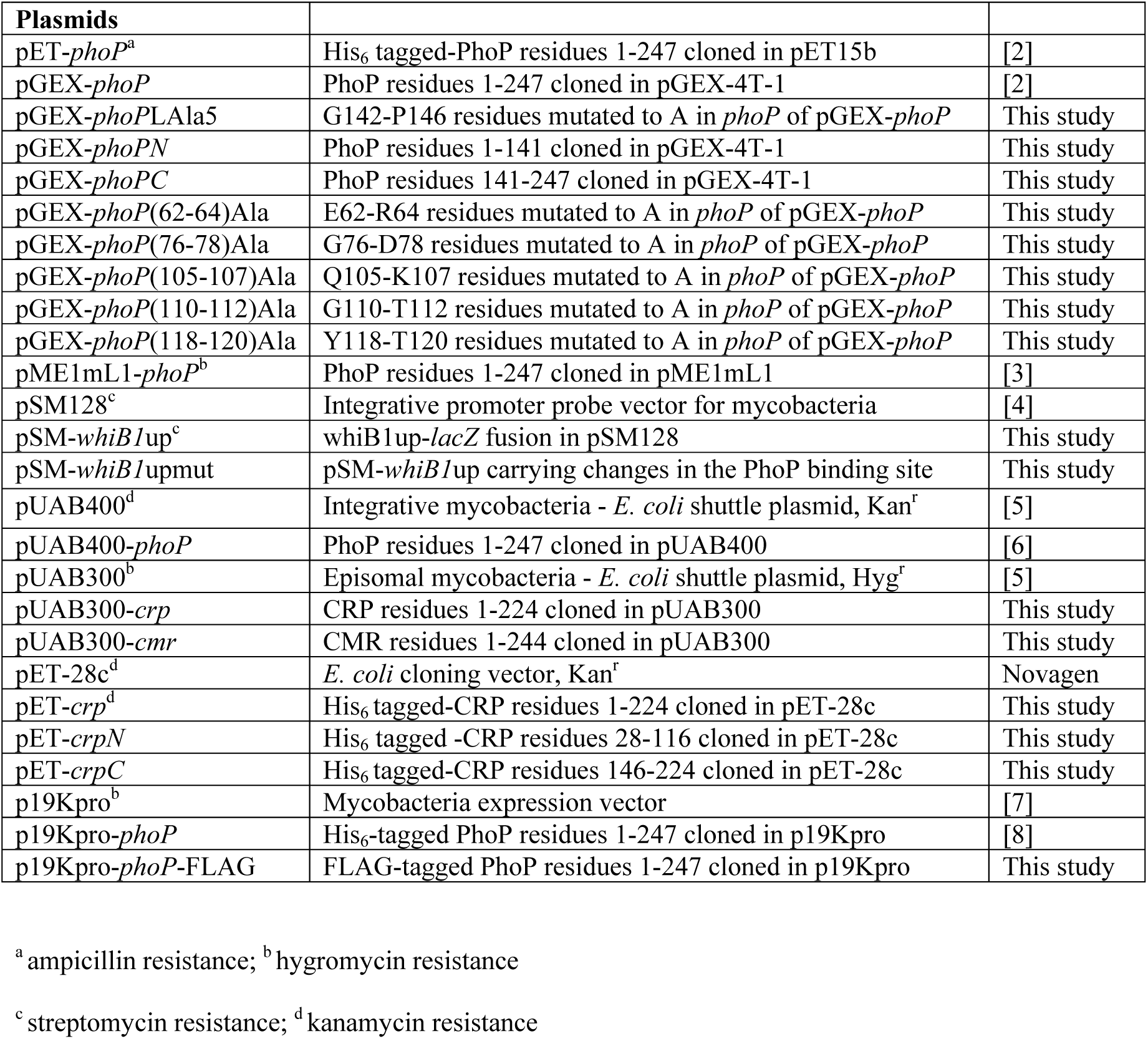
Plasmids used in this study

**Table S3.**
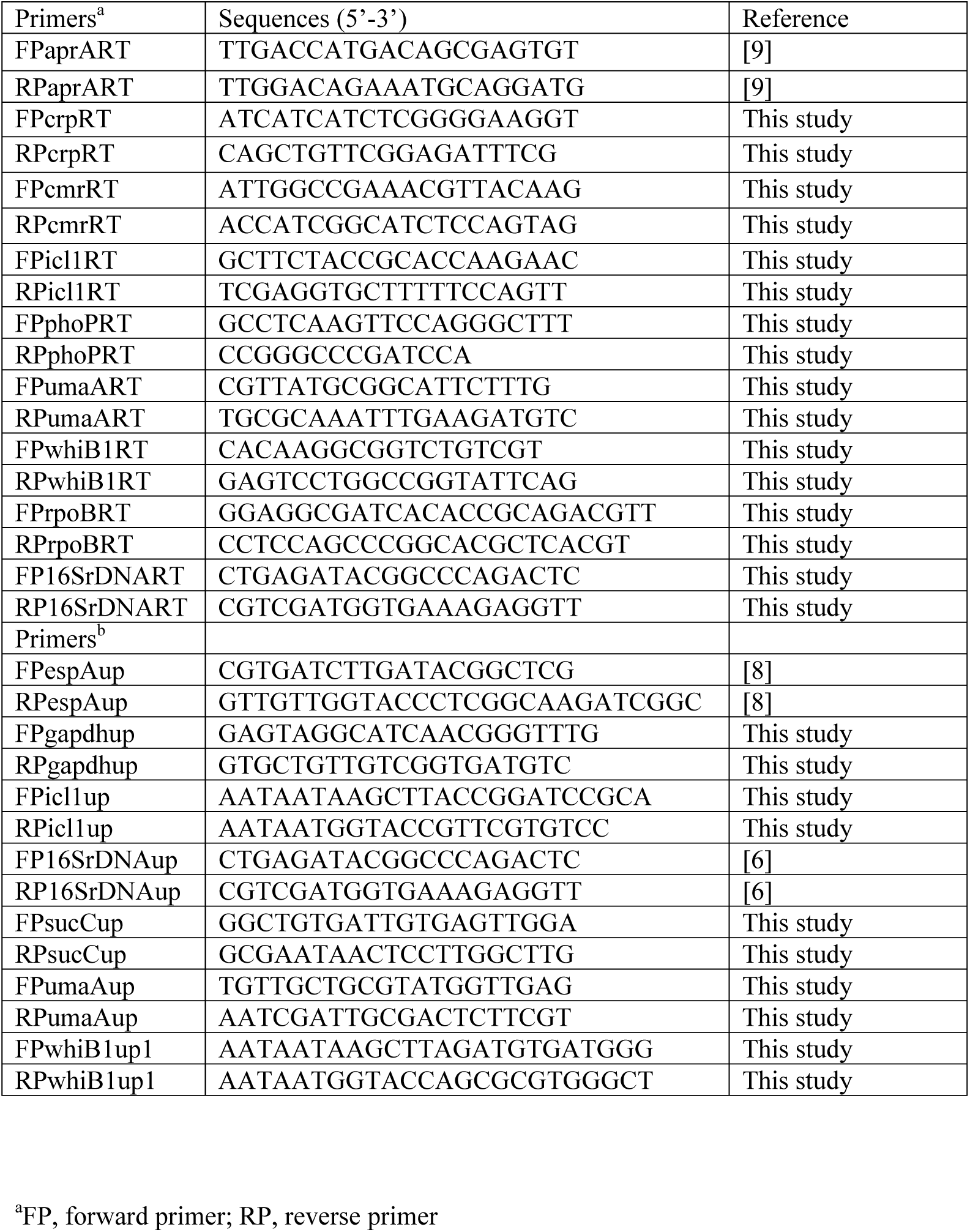
Sequence of oligonucleotide primers used in ^a^RT-qPCR and ^b^ChIP-qPCR experiments reported in this study

**Fig. S1:**
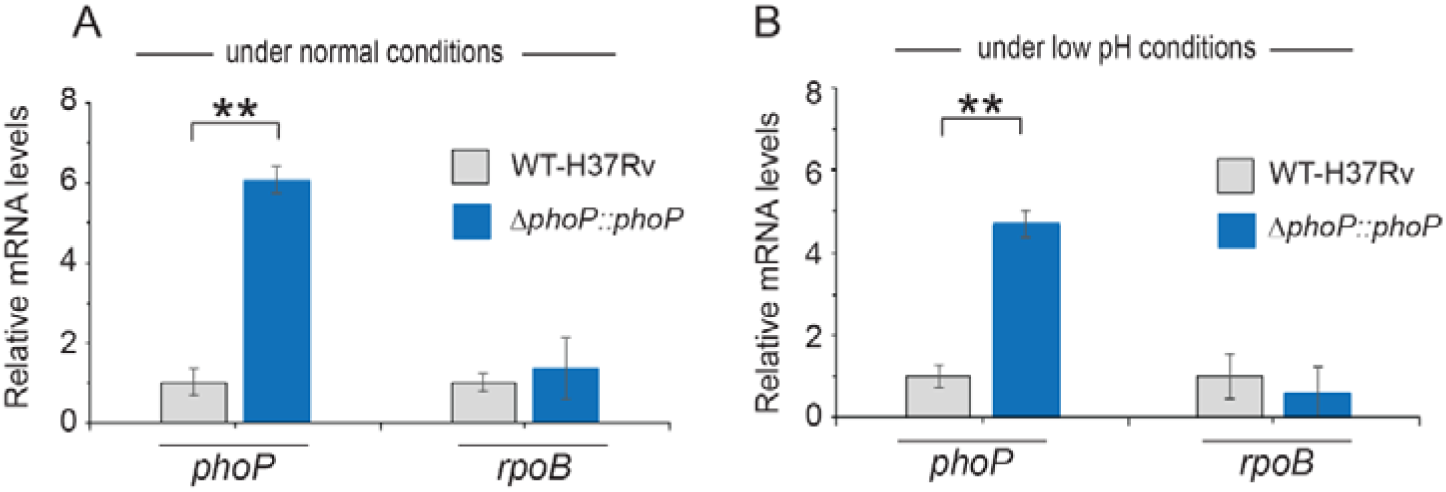
p*h*oP expression in WT-H37Rv and complemented *ΔphoP-*H37Rv under normal and acidic conditions of growth. Expression levels of *phoP* were measured in indicated mycobacterial strains grown under (A) normal pH (pH 7.0) and (B) acidic pH (pH 4.5) using RT-qPCR as described in the Methods. *rpoB* was used as a control, and the average fold differences were determined from two biological repeats (each with a technical repeat) (**P≤0.01).

**Fig. S2:**
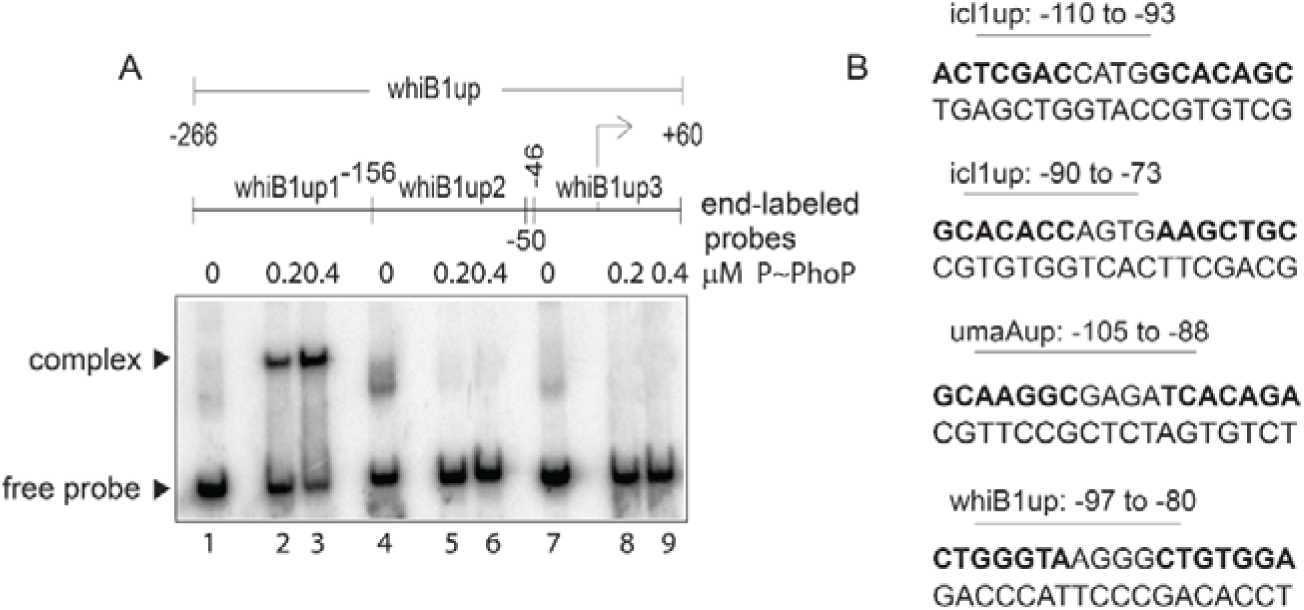
Probing core PhoP binding site within *whiB1* promoter region. (A) EMSA of radio-labelled whiB1up-derived fragments (whiB1up1-whiB1up3, as indicated) were performed with 0.2, and 0.4 µM of P∼PhoP (lanes 2-3, 5-6, and 8-9, respectively), pre-incubated in a phosphorylation mix with acetyl phosphate (AcP) as the phospho-donor, to probe core binding site of the regulator. The assay conditions, sample analyses, and detection of radio-labelled samples are described in the Methods; free probe and a slower moving complex are indicated on the figure. (B) Nucleotide sequences of likely PhoP binding sites within indicated promoters. The sequences are numbered relative to their corresponding transcription start sites.

**Fig. S3:**
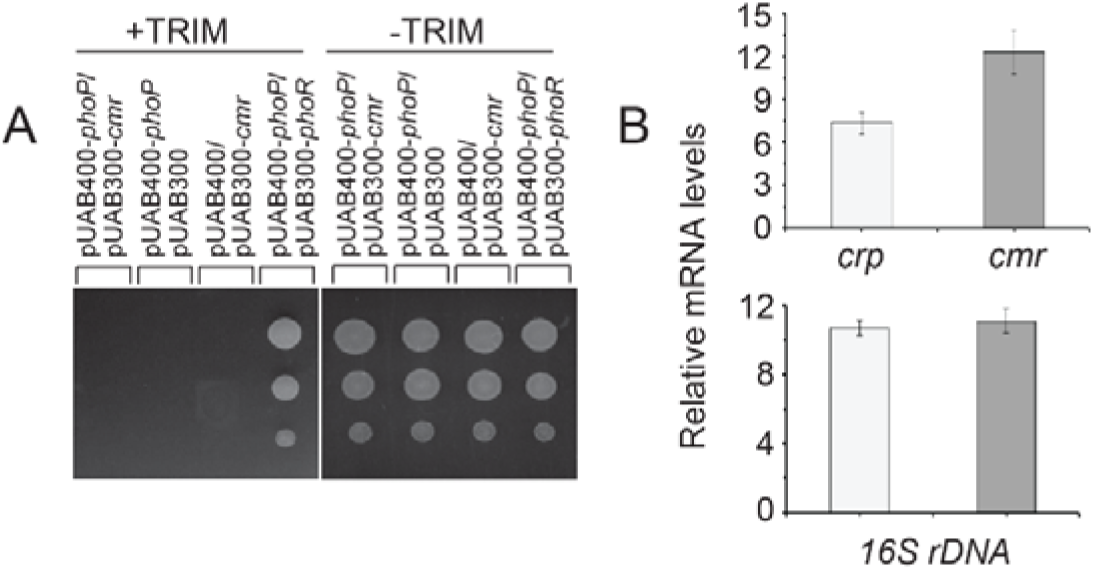
Probing PhoP-CMR interactions. (A) To probe protein-protein interaction by M-PFC assays, *M. smegmatis* co-expressing *M. tuberculosis* PhoP and cyclic AMP macrophage regulator (CMR), were grown on 7H10/hyg/kan plates in absence and presence of TRIM, and growth was examined. In both cases, empty vectors were included as negative controls, and co-expression of pUAB400-*phoP*/pUAB300-*phoR* (as a positive control) encoding PhoP and PhoR, respectively, showed *M. smegmatis* growth in presence of TRIM. All the strains grew well in absence of TRIM. (B) To verify expression of *M. tuberculosis* CRP and CMR in *M. smegmatis*, mRNA levels of the regulators were compared by RT-qPCR. Average fold changes in mRNA levels from two biological repeats (each with two technical repeats) are plotted, and non-significant difference is not indicated.

**Fig. S4:**
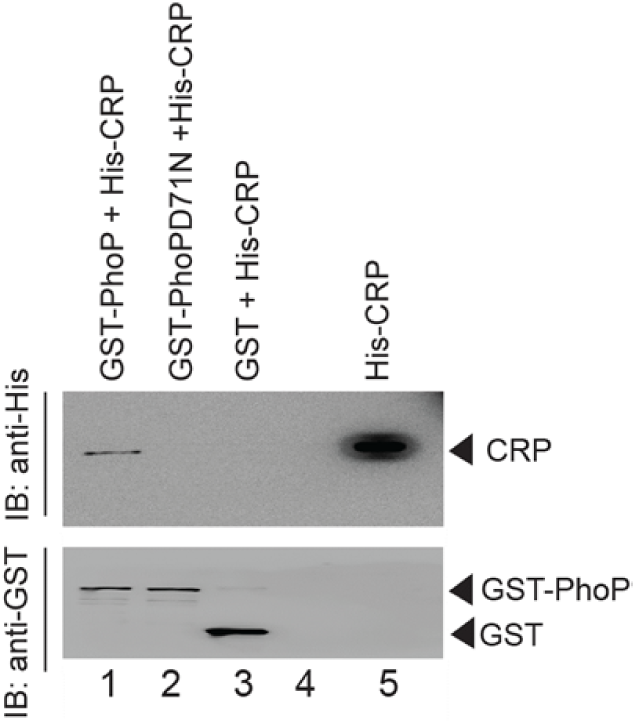
Phosphorylation-defective PhoP fails to interact with CRP. To examine whether phosphorylation of PhoP impacts CRP-PhoP interaction, crude lysates of cells expressing His_6_-tagged CRP was incubated with glutathione-Sepharose previously immobilized with GST-tagged PhoP (lane 1) or PhoPD71N (lane 2), carrying a single substitution of Asp-71 to Asn-71 and therefore, remains ineffective for phosphorylation at Asp-71. Analysis of bound fractions (lanes 1-2) was carried out as described in the legend to Fig. 5 and control sets include glutathione Sepharose immobilized with GST alone (lane 3), or the resin alone (lane 4); lane 5 resolved recombinant His_6_-tagged CRP.

**Fig. S5:**
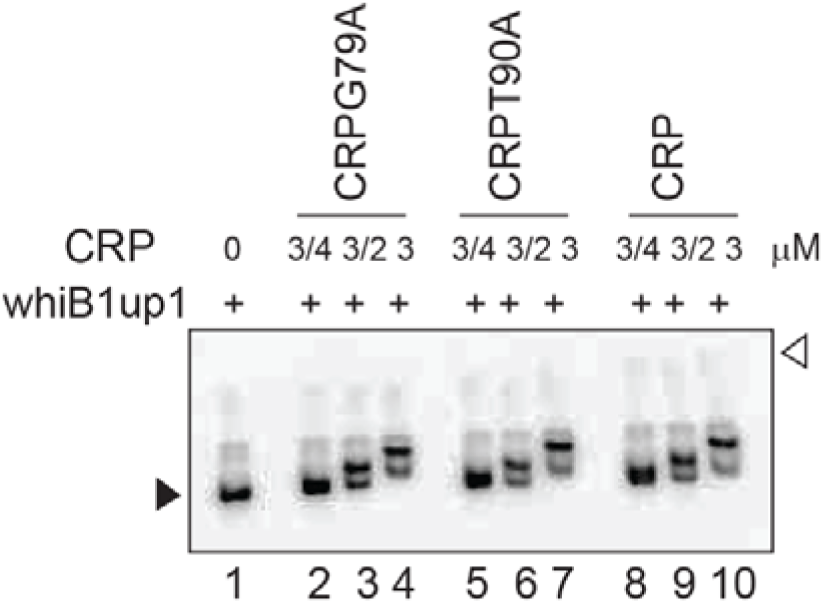
cAMP binding-defective CRP proteins are functional. To examine functionality of the mutant CRP proteins (CRPG79A, and CRPT90A), deficient for cAMP binding, EMSA of radio-labelled whiB1up was carried out using indicated mutants (lanes 2-7) and WT-CRP (as a positive control; lanes 8-10), respectively. The assay conditions, sample analyses, and detection of radio-active samples are described in the Methods. The filled and empty arrowheads on the left and right indicate free probe and the origin of the gel, respectively.

## Notes

### Competing Interest Statement

The authors have declared no competing interest.

